# Lower perceived stress enhances neural synchrony in perceptual and attentional cortices during naturalistic processing

**DOI:** 10.1101/2024.09.13.612956

**Authors:** J. Craig, C. Matisz, K. Klamer, C. Haines, K. Sullivan, C. Ekstrand

**Affiliations:** Ekstrand Neuroimaging Lab, Department of Neuroscience, University of Lethbridge 4401 University Dr W, Lethbridge AB, Canada, T1K 6T5

**Author notes:** Correspondence to: Chelsea Ekstrand, 4401 University Dr W, Lethbridge, AB, Canada, T1K 3M4.

**Keywords:** functional magnetic resonance imaging, perceived stress, cognitive processing, naturalistic cognition, intersubject correlation

## Abstract

Perceived stress is the subjective appraisal of the level of stress experienced by an individual in response to external or internal demands. Recent research on perceived stress has highlighted its role in influencing cognition, leading to a disruption in cognitive processes, such as emotional processing, attention, and perception. However, most neuroimaging studies examining stress have used static stimuli (e.g., still images) that do not encapsulate real-life multimodal processing in the brain. The current research uses data from the Naturalistic Neuroimaging Database (v2.0; Aliko et al., 2020) to examine differences in neural synchrony (as measured by intersubject correlations; ISCs) associated with perceived stress using functional magnetic resonance imaging (fMRI). We evaluated how self-reported perceived stress levels influence neural synchrony patterns in response to different naturalistic stimuli by examining the differences in neural synchrony between individuals with low and high perceived stress levels. We determined that lower perceived stress was observed with greater neural synchrony areas associated with perceptual and attention processing, including the lateral occipital cortex, superior temporal gyrus, superior parietal lobule, orbital frontal cortex, and the occipital pole. These results indicate that high levels of perceived stress heavily alter neural processing of complex audiovisual stimuli. Together, these results provide evidence that perceived stress influences cognitive processing in everyday life.

---

Stressful events produce both objective and subjective responses. Objective responses include physiological responses related to arousal, such as increased respiration and heart rate (Higgins, 2017), whereas subjective responses encompass thoughts, feelings, and emotions related to the experience (Wearne et al., 2019). This subjective stress response affects how individuals perceive and respond to stressful events in everyday life. Closely related to the concept of subjective stress is perceived stress, which is defined as the feeling or thoughts an individual has about how much stress they are under at a given time (Cohen et al., 1983). Perceived stress and objective stress responses are inherently correlated, as appraisal of external stressors (perceived stress) directly influences and shapes subsequent emotional and physiological responses (Aggarwal et al., 2014; Christensen et al., 2023; Xu et al., 2020). Stimulation of the stress system leads to alterations in behavior and physiological responses, enabling individuals to adjust and cope (Johnson et al., 1992). However, an exaggerated subjective response to stress has been demonstrated to increase the risk of adverse health outcomes, inhibit brain functions and decision-making processes, and lead to deficits in cognitive function that can negatively impact quality of life (Marin et al., 2011). Thus, perceived stress plays a pivotal role in the stress response that may compromise cognitive function (Phibbs et al., 2019).

Previous research suggests that heightened levels of perceived stress exert a profound impact on various cognitive functions such as attention, memory and emotional processing (Christensen et al., 2023; Higgins, 2017). For example, Chen et al. (2019) observed that higher levels of perceived stress predict poorer performance in multiple cognitive domains, including episodic memory, perceptual speed, and attention. Further, elevated stress levels diminish cognitive flexibility while promoting heightened salience for threat-related information, inhibiting overall cognitive performance (Christensen et al., 2023). Results from Liu et al. (2020) suggest that perceived stress also affects attention by reducing the amount of attentional resources available to allocate to incoming information. In line with these cognitive changes, heightened perceived stress is also associated with differences in brain structure and function, particularly in the prefrontal cortex, orbitofrontal cortex (OFC), insula, amygdala, and hippocampus, as well as visual and attention related brain areas (including the superior parietal lobule (SPL), lateral occipital cortex (LOC), and inferior parietal sulcus (IPS); Aggarwal et al., 2014; Fassett-Carman et al., 2022; Marin et al., 2011; Shackman et al., 2011; Zhu et al., 2022).

Empirical evidence suggests that heightened perceived stress can both enhance and disrupt perceptual processing depending on the magnitude. For example, low-to-moderate levels of stress have been suggested to improve information processing by enhancing attention and cognitive engagement (Shackman et al., 2011; Wang et al., 2023). However, excessive or prolonged psychological stress can lead to perceptual distortions and disruptions in attention and perceptual processing (Yaribeygi et al., 2017). Indeed, Tiferet-Dweck et al. (2016) proposed that stress and perceptual load compete for attentional resources, affecting the selective processing and conscious perception of stimuli. In line with this, Rojas-Thomas et al. (2023) observed that individuals with high stress levels exhibit disruptions in bottom-up and top-down attentional control. Further, stress-related changes in the prefrontal cortex (which is associated with attentional control and decision-making) can impair the filtering of information (Arnsten, 2009). Thus, perceived stress impacts how the brain processes incoming stimuli, however, the neural mechanisms related to these perceptual changes remains relatively unexplored.

It is important to note, however, that our knowledge of how perceived stress affects neural processing is dominated by studies that employ stress induction paradigms in laboratory settings (Berretz et al., 2021). While this research has contributed to our understanding of the role perceived stress plays in cognition, it often does not capture the circumstances in which stress is encountered within natural settings. In contrast, using naturalistic stimuli, such as films, music, and social interactions to study the stress response can enhance ecological validity by reflecting more “real-world” conditions (Sonkusare et al., 2019). Previous studies demonstrate that as stimuli become more naturalistic, there is a widespread increase in shared brain activation in regions associated with higher-order cognitive functions, context integration, and emotional processing, as naturalistic stimuli promote dynamic multimodal processing (Finn et al., 2018; Hasson, 2004; Sonkusare et al., 2019). Thus, naturalistic stimuli provide a rich and dynamic context for studying individual differences in cognitive and emotional processing. A common analytic approach that is used with naturalistic stimuli is known as intersubject correlation (ISC) analysis. ISC is a model-free approach that correlates neural responses across participants throughout the duration of a stimulus (Hasson, 2004) to identify shared neural responses across different experimental conditions by identifying voxels that consistently respond to a stimulus across subjects (Nastase et al., 2019).

Importantly, using naturalistic stimuli can allow researchers to observe how trait-level differences manifest in real-world settings, providing deeper insights into the underlying neural mechanisms associated with specific traits. For example, Finn et al. (2018) examined differences in neural synchrony between participants with either low or high levels of trait paranoia to an ambiguous auditory narrative. Results from this study showed that trait paranoia acted as an implicit prime for processing the naturalistic stimulus, resulting in unique patterns of brain synchrony for the high paranoia group in comparison to the low paranoia group. Additionally, Jajcay et al. (2024) identified differences in dynamic brain activity during naturalistic viewing can predict certain personality traits, such as neuroticism. Thus, naturalistic stimuli are a valuable tool for examining neural synchrony associated with specific cognitive and personality traits. However, to our knowledge, naturalistic stimuli have not been used to identify trait level differences in neural synchrony associated with perceived stress.

Based on this, our study aims to investigate the impact of perceived stress on the processing of naturalistic stimuli using an ISC paradigm. This will allow us to identify commonalities in neural processing using more “true-to-life” stimuli, providing valuable insight into perceptual changes associated with heightened perceived stress. To achieve this, we used data from the Naturalistic Neuroimaging Database (NNDb; version 2.0.0; Aliko et al., 2021) from participants who watched full-length audiovisual films during fMRI. We split participants into two groups based on their perceived stress scores (i.e., low-stress and high-stress) and analyzed the data using ISC analysis to identify patterns of neural synchrony for each group. Drawing from prior research, we hypothesized that lower scores on the PSS would be linked to greater synchrony in regions responsible for perception and attention, particularly in the primary and secondary visual and auditory cortices. This is due to the fact that higher stress levels have been found to disrupt these cognitive functions (Liu et al., 2020; Sandi, 2013; Wang et al., 2023). We expect that lower perceived stress scores would be associated with greater ISCs in areas such as the prefrontal cortex, OFC and, insula, as well as visual and attention related brain areas (including the, LOC and, attention areas, such as the SPL and IPS). By examining responses across a diverse set of films, we aim to identify generalizable patterns of neural processing that are influenced by perceived stress, rather than responses specific to a single stimulus. These findings will contribute to our understanding of how perceived stress impacts the processing of naturalistic stimuli, demonstrating that perceived stress alters general perceptual and cognitive processing in more true-to-life contexts. These findings will contribute to our understanding of how perceived stress impacts the processing of naturalistic stimuli.

## Methods

### Participants and Data

This study used preprocessed fMRI data from the NNDb (v2.0; Aliko et al., 2020) downloaded from Open Neuro (https://openneuro.org/datasets/ds002837/versions/2.0.0) from 86 participants (42 males/44 females, ages 19-53, mean age of 28.5 years) who underwent fMRI while viewing one of 10 full-length feature films. We chose to use all 10 films available from the NNDb in this study to ensure a comprehensive analysis that would not be limited to a single genre. By including a diverse range of films spanning multiple genres, we ensured that any differences in ISC between the low and high stress groups was not due to the specific choice of stimulus. This approach aligns with the methodology described in Aliko et al. (2020), which emphasizes the importance of diverse film selection to avoid bias towards a particular type of emotional response or genre-specific stimuli and instead aims to look at naturalistic stimuli in general. These 10 highly rated films spanned various genres from comedy to drama (for more information about how films were selected, see Aliko et al., 2020). All participants were right-handed, had normal or corrected-to-normal vision, did not have hearing impairments, were native English speakers, and had no history of neurological or psychiatric disorders (full details can be found in Aliko et al., 2020). Collection of the original data was approved by the ethics committee of University College London and participants provided written informed consent to participate in the study and share anonymized data. All methods were carried out in accordance with relevant guidelines and regulations (Aliko, 2020).

Data was acquired by Aliko et al. (2020) and full data acquisition details can be found in Aliko et al. (2020). Briefly, data was acquired on a 1.5T Siemens Magnetom Avanto MRI scanner using a 32-channel head coil. FMRI data was acquired using multiband echoplanar imaging (EPI) sequence with the following scanning parameters: repetition time (TR) of 1.0 s, echo time (TE) of 54 ms, flip angle of 75°, 40 interleaved slices, resolution 3.2 mm isotropic, with a 4x multiband factor with no in-plane acceleration. A T1-Magnetization Prepared Rapid Acquisition Gradient Echo (MPRAGE) anatomical scan was also acquired for each participant, with a TR of 2.73 s, TE of 3.57 ms, and a resolution of 1.0 mm³. Noise-attenuating headphones were used to ensure that audio quality was clear. The films were shown through a mirror-reversed LCD projector viewed through a mirror mounted on the head coil and played with minimal interruptions. Participant attentiveness was monitored via a camera focused on their eyes. Due to the constraints of the EPI sequence and software, the films were presented in segments lasting 40–50 minutes each. These breaks were deliberately timed to coincide with scenes that did not contain vital plot information or dialogue, ensuring minimal interference with the participant experience.

Data preprocessing was conducted by Aliko et al. (2020). First, the separate runs for each session were concatenated across time with a timing correction using AFNI’s ’3dTproject’ (Cox,1996; Taylor et al., 2018). Next, all functional scans underwent a comprehensive set of preprocessing steps. These steps included time shifting (aligning the data with the timing of the stimulus), despiking (the removal of extreme and sudden signal fluctuations), volume registration (spatially aligning the acquired volumes), MNI alignment (standardizing all functional data to a template), masking the time-series, smoothing with a 6 mm full width half max kernel, detrending with regressors (e.g., to address motion artifacts), additional timing correction, and manual ICA denoising. These functions were done by employing AFNI’s afni_proc.py pipeline (Cox, 1996; Taylor et al., 2018). For this analysis, we used preprocessed functional files that had undergone spatial smoothing, while retaining their original content without any censoring (i.e., “uncensored” files).

### Behavioural Questionnaires

Once the MRI imaging session was completed, participants were asked to complete all questionnaires from the National Institute of Health (NIH) Toolbox, which includes 23 validated comprehensive questionnaires that measures of sensory, motor, cognitive, and emotional processing, as well as identify specific traits to assess individual differences (Aliko, 2021; Salsman et al., 2013). In the present study, we specifically used the PSS raw scores quantified using the NIH Toolbox 2.0 Perceived Stress Scale (PSS) questionnaire done by participants (see Figure 1 for the distribution of PSS scores). The NIH Toolbox PSS questionnaire evaluates psychological responses to stress over a month-long period (Cohen et al., 1983). To determine if there were any correlations between the PSS scores and other trait scores within the NIH Toolbox, we used SPSS (version 27; IBM Corp., 2020) to run a bivariate correlation between PSS scores and all other questionnaires. To observe if there were any differences between in age between the low and high PSS groups, we used SPSS (version 27; IBM Corp., 2020) to run an independent samples *t-*test.

**Figure 1.**
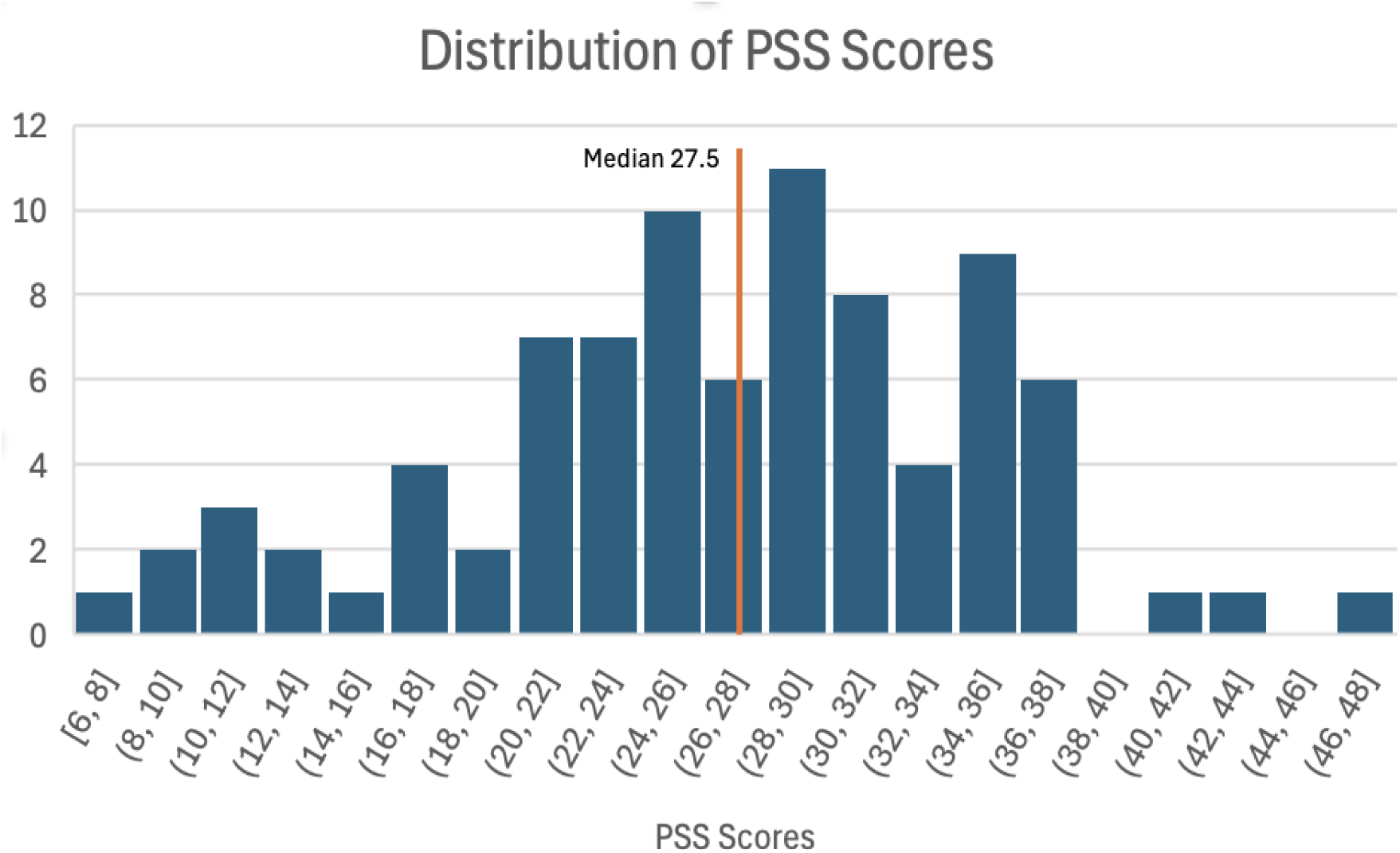
The distribution of T-scores from the NIH Toolbox 2.0 Perceived Stress Scale questionnaire for all participants N=86 who viewed various films within the Naturalistic Neuroimaging Database v2.0. Range 6-48, median 27.5.

### Data Analysis

#### Defining low and high stress groups

To separate our participants into low and high stress groups, we used a median split on all participants Perceived Stress Fixed Form Age 18+ v2.0 scores from the NIH Toolbox (Salsman et al., 2013; range 6-48, median = 27.5, standard deviation = 8.11). This resulted in 43 participants in the low-stress group (21 females, mean age = 29, standard deviation 3.7) and 43 participants in the high-stress group (19 females, mean age = 27, standard deviation 4.5). For the low stress group, the median PSS score was 22 and the standard deviation was 5.6, For the high stress group, the PSS median score was 33 with a standard deviation of 4.2.

#### Intersubject Correlation Analysis

To perform ISC analysis across all possible participant pairs in all films, we modified the number of volumes in 9 of the 10 films to align with the number of volumes associated with the shortest film, a total of 5470 volumes, using *fslroi* (Jenkinson et al., 2012). We then performed ISC analysis across all participants both within and between movie conditions. In the present study, ISCs were computed for all unique pairs of participants both within- and between-perceived stress groups, which produced 3655 (*n*\*(*n*-1)/2, where n = 86) unique ISC maps. This resulted in a total of 903 unique participant pairings for the low-stress group, and 903 unique pairings for the high-stress group (i.e., 903 Low-low stress pairs and 903 High-high stress pairs) and 3655 between group pairings (i.e., Low-high stress pairs). Importantly, the number of within movie pairs (i.e., participants who watched the same film) and between movie pairs (i.e., participants who watched different films) was equal for each group, thus higher ISC in one group should not be due to overrepresentation of ISCs from a single film.

#### Linear mixed-effects modeling

To identify significant ISC for each contrast, we used linear mixed effects (LME) modelling, implemented via the *3dISC* module in AFNI (Cox, 1996; as described by Chen et al., 2020). This approach identifies significant regions of neural synchrony for different contrasts across participants, detecting brain areas that respond similarly to the stimulus. Further, LME accounts for the complex covariance structure of ISC data by incorporating both fixed and random effects within a unified model and has been shown to accurately control for false positives in ISC analysis (Chen, 2017). LME assumes the response variable is normally distributed, and therefore we used the Fisher transformed pairwise ISCs for this analysis. The utility of LME models for modeling ISC data has been extensively evaluated by Chen and colleagues (G. Chen et al., 2016; 2017; 2019) and has been shown to be efficient and robust, even in comparison to non-parametric analysis methods.

We modelled the following contrasts: average ISC in the low-stress group (Low-low stress), average ISC in the high-stress group (High-high stress), average ISC between the low and high groups (Low-high stress), comparison of ISC between the low- and high-stress groups (Low-low vs. High-high stress contrast), comparison of ISC between the low-stress group and between group ISC (Low-low vs. Low-high stress contrast), and comparison of ISC between the between group ISC and high-stress group (Low-high vs. High-high stress contrast). Significant ISC was determined using a voxel-wise false discovery rate (FDR) threshold of *q* < 0.001 for our within-group average ISCs (i.e., Low-low perceived stress, High-high perceived stress), and a threshold of *q* < 0.05 for our between-group contrasts (i.e., Low-low vs. High-high perceived stress, Low-low vs. Low-high perceived stress, Low-high vs. High-high perceived stress).

Additionally, to ensure that our results were not due to closeness of scores (i.e., participants with more similar scores showing similar brain activation, regardless of the magnitude of the perceived stress score) we calculated the absolute difference between each pair of participants PSS scores and used this value as a predictor in an LME model for pairwise ISC (abs(PSS(pairwise similarity)) + (1|sub1) + (1|sub2)). To do so, we subtracted the PSS score of the second participant in each pair from the PSS score of the first participant, took the absolute value, and included this number in the new LME model. This approach allows us to determine if the ISC results are due to the similarity in PSS scores, rather than an association between higher ISC and either lower or higher PSS scores (Finn et al., 2018; Jangraw et al., 2023). We then applied an FDR correction to the resulting *p*-values to identify significant ISCs. Significant ISCs for the group comparison contrasts were defined using a voxel wise false discovery rate (FDR) of *q* < 0.05. Significant results were transformed into surface space for visualization purposes only.

## Results

### Behavioral Questionnaires

We used SPSS (version 27; IBM Corp., 2020) to perform a bivariate correlation among the different NIH toolbox dimensions and PSS score to determine which other questionnaires were significantly correlated with the PSS. Significant correlations were identified at a *p*-value of .001. The following questionnaires were significantly correlated at the *p <* 0.001 level: Loneliness (*r*^2^ = 0.321, *p* = 0.022), Anger Affect (*r*^2^ = 0.342, *p* = 0.031), and Anger Hostility (*r*^2^ = 0.642, *p* = 0.001). The following questionnaires were negatively correlated at the *p <* 0.001 level: Positive Affect (*r*^2^ = -0.532, *p* = 0.001), General Life Satisfaction (*r*^2^ = -0.541, *p* = 0.001), Friendship (*r*^2^ = -.401, *p* = 0.001), Self-Efficacy (*r*^2^ = -.491, *p* = 0.001), Meaning and Purpose (*r*^2^ = -0.422, *p* = 0.001), and Instrumental Support (*r*^2^ = -0.221 *p* =0.041). We also conducted an independent samples t-test to determine if there was a significant difference in age between the low- and high-stress groups. We did not find a significant difference in age between the low- and high-stress groups, *t* (84) = 0.623, *p* = 0.611.

### Low-low and High-high Stress Group Averages

Results for the Low-low group average contrast can be found in Figure 2 and Supplementary Materials A Table S2. The LME contrast for the low-stress ISC group average (i.e., Low-low stress) revealed significant neural synchrony across frontal, occipital, and parietal cortices. Significant areas of synchrony included the bilateral posterior cingulate cortex (PCC), bilateral temporal parietal junction (TPJ), and bilateral orbitofrontal cortex (OFC). Other regions that showed significant neural synchrony were the bilateral supramarginal gyrus, parietal operculum, occipital cortex, calcarine sulcus, lingual gyrus, precuneus and temporo-occipital cortices.

**Figure 2:**
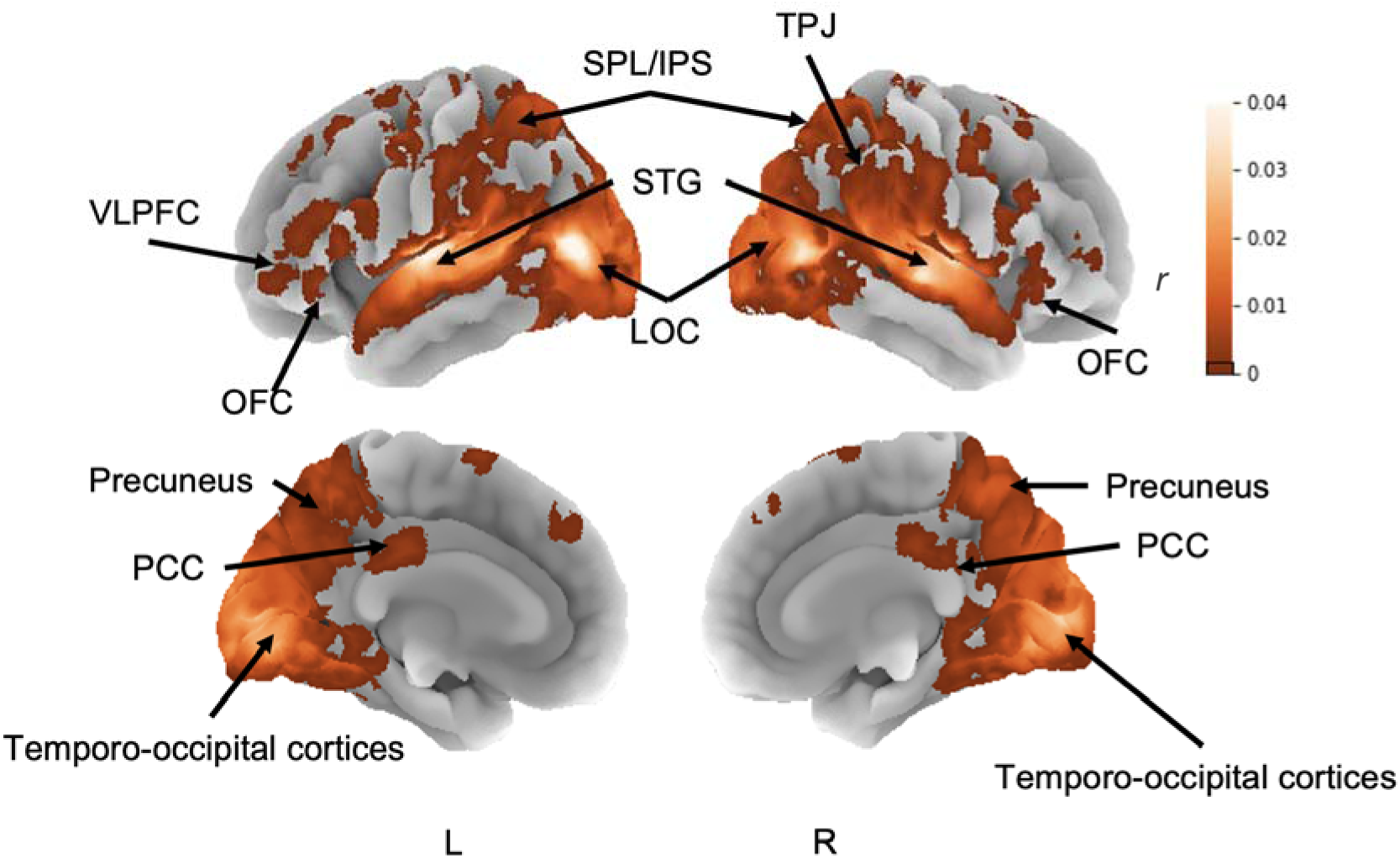
Significant ISCs from the low-low stress group average contrast. Results are displayed as an FDR set to *q* < 0.001 and only show clusters greater than 5 voxels. Maps are shown in the lateral and medial orientations.

Average neural synchrony for the high stress group ISC average (i.e., High-high stress) as seen in Figure 3 and in Supplementary Materials A Table S3, showed significant areas of synchrony in the bilateral parahippocampal gyrus (PHG), inferior frontal gyrus (IFG), LOC, parietal operculum, lingual gyrus, intracalcarine cortex, superior temporal gyrus (STG), parietal operculum, right insula, and left SPL. Widespread neural synchrony across the temporal, occipital, parietal, and frontal lobes (primarily in primary and secondary visual and auditory areas) is in line with previous research (Güçlütürk et al., 2018; Hasson, 2004) demonstrating that widespread synchrony results from watching an audiovisual stimulus.

**Figure 3.**
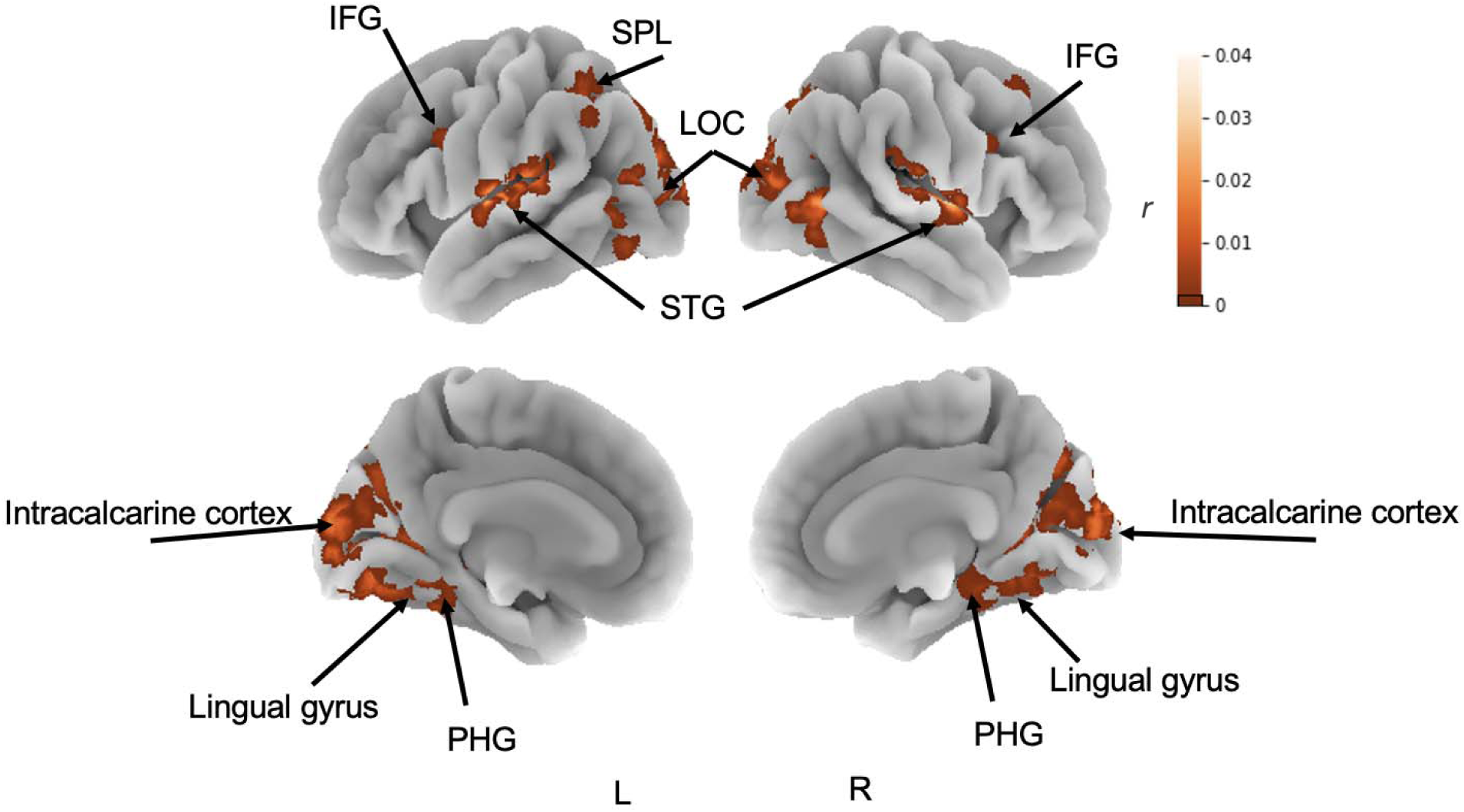
Significant ISCs from the High-stress group average contrast. Results are displayed as an FDR set to *q* < 0.001 and only show clusters of 5 or more voxels. Maps are shown, in the lateral and medial orientations.

### Low-low vs. High-high Stress Contrasts

#### Low-low > High-high stress

Significant neural synchrony for Low-low > High-high contrast can be found in Figure 4 (warm colours) and Supplementary Materials A Table S4. We observed significant ISC in the right OFC, bilateral LOC, bilateral STG, bilateral occipital pole, and bilateral SPL.

**Figure 4:**
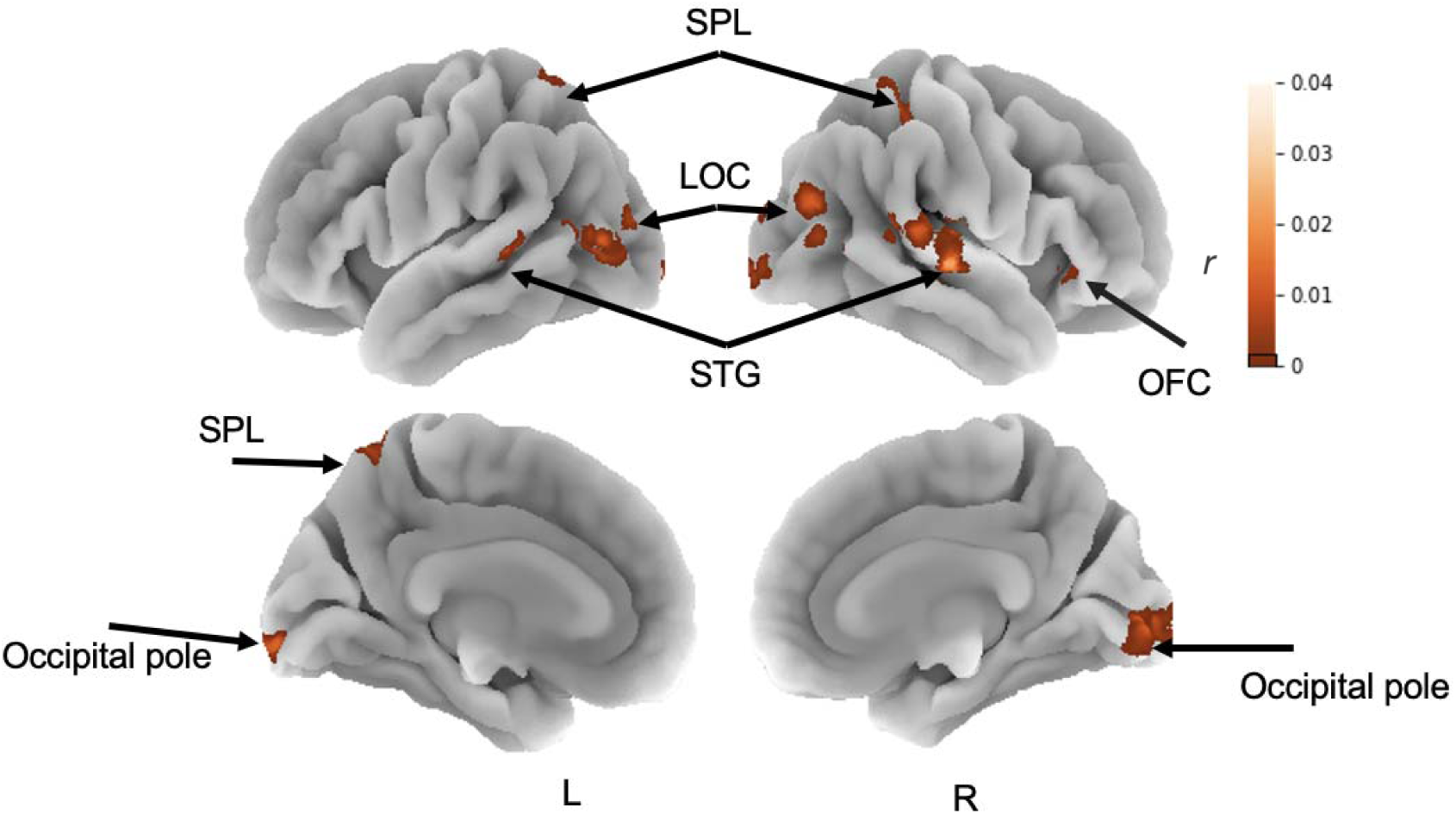
Significant ISCs from the Low-low vs. High-high stress contrast. Results for Low-low > High-high are shown in warm colours and results for High-high > Low-low are shown in cold colours. Results are displayed with an FDR set to *q* < 0.05 and only show clusters of 5 or more voxels. Maps are shown for the left (L) and Right (R) hemispheres in the lateral and medial orientations.

#### High-high > Low-low stress

There were no significant ISCs for this contrast.

#### Low-low and High-high vs. Low-High stress

We found no significant ISCs when comparing ISC within the low-stress group to between-group ISC (low-low vs. low-high stress contrast). Similarly, there were no significant ISCs when comparing ISC within the high-stress group to between-group ISC (low-high vs. high-high stress contrast).

### Absolute Distance of PSS Scores

We calculated the absolute difference between each pair of participants PSS scores and used this measure as a predictor in our LME model to examine its effect and use as a predictor on pairwise ISC. We did not find any significant ISCs following the analysis as it was revealed that there was no significant relationship between the absolute differences in PSS scores and the ISC’s, suggesting that the similarity in stress levels does not account for the differences in ISC between low and high stress groups illustrated previously. Instead, the observed increase in ISC appears to be associated with lower individual PSS scores. This finding suggests that participants with lower perceived stress scores exhibit greater neural synchrony than participants with higher perceived stress scores.

## Discussion

This experiment sought to investigate how perceived stress impacts processing of naturalistic stimuli in the brain. To do so, we separated participants into low and high perceived stress groups based on their PSS scores and examined how subjective stress levels impacted neural synchrony towards different complex audiovisual films that varied in genre and emotional valence. Of particular importance, we investigated differences in brain synchrony between individuals with low and high perceived stress levels across a diverse array of 10 audiovisual movies. This comprehensive analysis provides critical insights into how perceived stress levels impact more real-world naturalistic processing, demonstrating the pervasive influence of stress on cognitive and neural functions, regardless of the specific stimuli involved. We also calculated the absolute distance between pairwise PSS scores to ensure that differences in ISC were not due to similarity in PSS score, regardless of the magnitude. Based on previous research, we hypothesized that low PSS scores would be associated with greater synchrony in perceptual (i.e., primary and secondary visual and auditory cortices) and attentional areas, as heightened stress levels have been shown to disrupt these processes (Liu et al., 2020; Rojas-Thomas et al., 2023; Shackman et al., 2011; Tiferet-Dweck et al., 2016). Our primary results are highlighted below.

Results from the low and high-stress group average ISC analyses showed widespread neural synchrony across the occipital, temporal, parietal, and frontal cortices, most notably in primary and secondary visual and auditory areas and relevant association cortices as a result of viewing the same audiovisual stimulus (Finn et al., 2018; Güçlütürk et al., 2018; Hasson, 2004; Jääskeläinen et al., 2021). This is in line with previous research showing widespread neural synchrony in response to audiovisual movie stimuli (e.g., Finn et al., 2018; Güçlütürk et al., 2018; Hasson, 2004; Jääskeläinen et al., 2021), and extends these findings across 10 different films.

When comparing participants with lower PSS scores to participants with higher stress scores, we found significant differences in neural synchrony in areas associated with auditory, visual, attention, and reward related processes. When examining differences between the low and high perceived stress groups, we found greater neural synchrony for the low stress group than the high stress group in the bilateral STG (involved in auditory processing; Bigler et al., 2007), bilateral occipital pole (involved in visual processing; Nagy et al., 2012), bilateral intraparietal sulcus/SPL (involved in attention, visuospatial attention and working memory; Alahmadi, 2021), and right OFC (involved in emotional processing and representation of emotional stimuli; Welborn et al., 2009). Notably, we observed significant synchrony in the right OFC, which is involved in motor inhibition, emotion regulation, reward-related processing, and response inhibition (Viviani, 2014; Yang et al., 2020). This is in line with previous findings which suggest that increases in stress can affect and disrupt emotional processing (Christensen et al., 2023; Higgins, 2017). Therefore, compared to individuals with higher levels of perceived stress, individuals with lower levels of perceived stress show increased synchrony in frontal cortices as well as primary and secondary auditory and visual cortices, as this could be due to stress affecting visual and auditory attention and perception (Tiferet-Dweck et al., 2016; Wang et al., 2023).

In line with our hypothesis and previous research, we observed significantly greater neural synchrony for the low stress group than the high stress group in areas involved in perception and attention, including the LOC, occipital pole, and SPL. Comparatively, it has been shown that heightened perceived stress affects attentional resources available to be allocated towards incoming information, and may impact the cognitive processing of incoming stimuli (Liu et al., 2020; Shackman et al., 2011). Of note, we also found increased synchrony in the right OFC (involved in reward-related processing; Noonan et al., 2011; Rudebeck & Rich, 2018) for the low-stress group. The right OFC plays a key role in decision-making, emotional regulation, and reward processing (Welborn et al., 2009), and the STG, occipital pole, and LOC are essential for auditory and visual processing, visual perception, and object recognition (Catani et al., 2003; Nagy et al., 2012). This suggests that individuals with lower stress show more similar processing in regions associated with reward sensitivity, auditory, and visual processing compared to those with higher perceived stress.

In contrast, we did not find any regions that showed greater synchrony for the high stress group than the low stress group, suggesting high perceived stress is associated with more variable neural processing. This is in line with previous research showing that high perceived stress leads to more variable cognitive processing in multiple domains, including episodic memory, perceptual speed, and attention (Archer et al., 2017; Colibazzi et al., 2010; Shackman et al., 2011). This is further corroborated by the results from our LME analysis that used absolute distance in PSS scores as an explanatory variable to explore whether similarity of stress scores, rather than the magnitude of stress scores, could be driving correlations. To do so, we subtracted the PSS score of one participant in each pair from the other, took the absolute value of this difference, and included it in the LME model as a predictor for pairwise ISC. Results from this analysis did not show any significant ISCs, suggesting that the similarity in PSS scores is not driving heightened ISC for the low stress group. Ultimately, these results demonstrate the unique modulatory affect lower perceived stress has on cognitive processes involved in everyday life.

There are several limitations to this study. Because we used data from an online database, we did not have control over the stimuli that were presented to the participants, limiting our control over plot content, length, and genre of the films chosen. In addition, the data used in these experiments was collected using a 1.5T MRI, which has lower spatial specificity and a lower signal-to-noise ratio than stronger MRIs. The use of a higher tesla scanner would better encapsulate finer details of functional activation. Finally, this study assessed stress levels using self-report measures, possibly introducing biases and inaccuracies, which importantly contained no measure of physiological stress responses. Future research should consider incorporating objective measures of stress, such as physiological markers (e.g., cortisol levels, heart rate, galvanic skin response), to complement self-report measures that would provide a more comprehensive understanding of stress levels in future experiments.

In conclusion, this research provides novel insight into how perceived stress impacts cognition using a diverse set of naturalistic audiovisual movie stimuli. Our findings demonstrate that lower levels of perceived stress are associated with increased neural synchrony in perceptual (i.e., primary and secondary visual and auditory cortices) and attentional regulatory areas such as the bilateral SPL (involved in visuospatial attention; Wu et al., 2016), right OFC (primarily involved in sensory integration and reward related decision making; Rudebeck & Rich, 2018), bilateral STG (Highly involved in auditory processing), bilateral occipital pole, and bilateral LOC, both of which are involved in higher order visual representations and processing (Rehman & Al Khalili, 2023). These results suggest that perceived stress may serve as an implicit prime that alters the processing of real-world stimuli, particularly in regions associated with visuospatial processing, reward related processing, attention, and cognition (Alahmadi, 2021; Aminoff et al., 2013; Hampshire et al., 2010; Nagy et al., 2012; Namkung et al., 2017). Understanding how perceived stress influences neural processing of real-world stimuli enhances our knowledge of the impact of psychological stress on cognition and emotional processing. These insights provide a deeper understanding of how stress shapes our perceptions and interactions in daily life, highlighting its complex effects on our experiences and responses to diverse emotional contexts.

## References

Aggarwal, N. T., Wilson, R. S., Beck, T. L., Rajan, K. B., Mendes de Leon, C. F., Evans, D. A., & Everson-Rose, S. A. (2014). Perceived Stress and Change in Cognitive Function Among Adults 65 Years and Older. Psychosomatic Medicine, 76(1), 80. 10.1097/PSY.0000000000000016

Alahmadi, A. A. S. (2021). Investigating the sub-regions of the superior parietal cortex using functional magnetic resonance imaging connectivity. Insights into Imaging, 12(1), 47. 10.1186/s13244-021-00993-9

Aliko. (2021). *Naturalistic Neuroimaging Database {Data Set}.* Openneuro. 10.18112/OPENNEURO.DS002837.V2.0.0

Aminoff, E. M., Kveraga, K., & Bar, M. (2013). The role of the parahippocampal cortex in cognition. Trends in Cognitive Sciences, 17(8), 379–390. 10.1016/j.tics.2013.06.009

Archer, J. A., Lee, A., Qiu, A., & Annabel Chen, S.-H. (2017). Functional connectivity of resting-state, working memory and inhibition networks in perceived stress. Neurobiology of Stress, 8, 186–201. 10.1016/j.ynstr.2017.01.002

Arnsten, A. F. (2009). Stress signalling pathways that impair prefrontal cortex structure and function. Nature Reviews Neuroscience, 11(7), 410–422.

Bigler, E. D., Mortensen, S., Neeley, E. S., Ozonoff, S., Krasny, L., Johnson, M., Lu, J., Provencal, S. L., McMahon, W., & Lainhart, J. E. (2007). Superior temporal gyrus, language function, and autism. Developmental Neuropsychology, 31(2), 217–238. 10.1080/87565640701190841

Catani, M., Jones, D. K., Donato, R., & ffytche, D. H. (2003). Occipito temporal connections in the human brain. Brain, 126(9), 2093–2107. 10.1093/brain/awg203

Chen, G. (2017). Untangling the Relatedness among Correlations, Part II: Inter-Subject Correlation Group Analysis through Linear Mixed-Effects Modeling. Neuroimage, 147, 825–840.

Chen, Y., Liang, Y., Zhang, W., Crawford, J. C., Sakel, K. L., & Dong, X. (2019). Perceived Stress and Cognitive Decline in Chinese-American Older Adults. Journal of the American Geriatrics Society, 67(S3), S519–S524. 10.1111/jgs.15606

Christensen, D. S., Garde, E., Siebner, H. R., & Mortensen, E. L. (2023). Midlife perceived stress is associated with cognitive decline across three decades. BMC Geriatrics, 23(1), 121. 10.1186/s12877-023-03848-8

Colibazzi, T., Posner, J., Wang, Z., Gorman, D., Gerber, A., Yu, S., Zhu, H., Kangarlu, A., Duan, Y., Russell, J. A., & Peterson, B. S. (2010). Neural systems subserving valence and arousal during the experience of induced emotions. *Emotion (Washington*, D.C.), 10(3), 377–389. 10.1037/a0018484

Fassett-Carman, A. N., Smolker, H., Hankin, B. L., Snyder, H. R., & Banich, M. T. (2022). Neuroanatomical Correlates of Perceived Stress Controllability in Adolescents and Emerging Adults. *Cognitive, Affective*, & Behavioral Neuroscience, 22(4), 655–671. 10.3758/s13415-022-00985-2

Finn, E. S., Corlett, P. R., Chen, G., Bandettini, P. A., & Constable, R. T. (2018). Trait paranoia shapes inter-subject synchrony in brain activity during an ambiguous social narrative. Nature Communications, 9(1), Article 1. 10.1038/s41467-018-04387-2

Geerligs, L., Rubinov, M., Cam-CAN, & Henson, R. N. (2015). State and Trait Components of Functional Connectivity: Individual Differences Vary with Mental State. The Journal of Neuroscience, 35(41), 13949–13961. 10.1523/JNEUROSCI.1324-15.2015

Güçlütürk, Y., Güçlü, U., van Gerven, M., & van Lier, R. (2018). Representations of naturalistic stimulus complexity in early and associative visual and auditory cortices. Scientific Reports, 8(1), Article 1. 10.1038/s41598-018-21636-y

Hampshire, A., Chamberlain, S. R., Monti, M. M., Duncan, J., & Owen, A. M. (2010). The role of the right inferior frontal gyrus: Inhibition and attentional control. NeuroImage, 50(3), 1313–1319. 10.1016/j.neuroimage.2009.12.109

Hasson, U. (2004). Intersubject synchronization of cortical activity during natural vision. Science, 303(5664), 1634–2640.

Higgins, G. (2017, August 28). The role of cognition in stress: Relationship between perceived life stress, self-efficacy, optimism, self-esteem, positive affect, negative affect, and work stress. https://www.semanticscholar.org/paper/The-role-of-cognition-in-stress%3A-Relationship-life-Higgins/002b9266e3382959276dcad6ac833f37a722a7b9

Jääskeläinen, I. P., Sams, M., Glerean, E., & Ahveninen, J. (2021). Movies and narratives as naturalistic stimuli in neuroimaging. NeuroImage, 224, 117445. 10.1016/j.neuroimage.2020.117445

Jangraw, D. C., Finn, E. S., Bandettini, P. A., Landi, N., Sun, H., Hoeft, F., Chen, G., Pugh, K. R., & Molfese, P. J. (2023). Inter-subject correlation during long narratives reveals widespread neural correlates of reading ability. NeuroImage, 282, 120390. 10.1016/j.neuroimage.2023.120390

Johnson, E. O., Kamilaris, T. C., Chrousos, G. P., & Gold, P. W. (1992). Mechanisms of stress: A dynamic overview of hormonal and behavioral homeostasis. Neuroscience & Biobehavioral Reviews, 16(2), 115–130. 10.1016/S0149-7634(05)80175-7

Liu, Q., Liu, Y., Leng, X., Han, J., Xia, F., & Chen, H. (2020). Impact of Chronic Stress on Attention Control: Evidence from Behavioral and Event-Related Potential Analyses. Neuroscience Bulletin, 36(11), 1395–1410. 10.1007/s12264-020-00549-9

Marin, M.-F., Lord, C., Andrews, J., Juster, R.-P., Sindi, S., Arsenault-Lapierre, G., Fiocco, A. J., & Lupien, S. J. (2011). Chronic stress, cognitive functioning and mental health. Neurobiology of Learning and Memory, 96(4), 583–595. 10.1016/j.nlm.2011.02.016

Nagy, K., Greenlee, M., & Kovács, G. (2012). The Lateral Occipital Cortex in the Face Perception Network: An Effective Connectivity Study. Frontiers in Psychology, 3. https://www.frontiersin.org/articles/10.3389/fpsyg.2012.00141

Namkung, H., Kim, S.-H., & Sawa, A. (2017). The insula: An underestimated brain area in clinical neuroscience, psychiatry, and neurology. Trends in Neurosciences, 40(4), 200–207. 10.1016/j.tins.2017.02.002

Noonan, M. P., Mars, R. B., & Rushworth, M. F. S. (2011). Distinct Roles of Three Frontal Cortical Areas in Reward-Guided Behavior. Journal of Neuroscience, 31(40), 14399–14412. 10.1523/JNEUROSCI.6456-10.2011

Phibbs, S., Stawski, R. S., MacDonald, S. W. S., Munoz, E., Smyth, J. M., & Sliwinski, M. J. (2019). The influence of social support and perceived stress on response time inconsistency. Aging & Mental Health, 23(2), 214–221. 10.1080/13607863.2017.1399339

Rehman, A., & Al Khalili, Y. (2023). Neuroanatomy, Occipital Lobe. In StatPearls. StatPearls Publishing. http://www.ncbi.nlm.nih.gov/books/NBK544320/

Rojas-Thomas, F., Artigas, C., Wainstein, G., Morales, J.-P., Arriagada, M., Soto, D., Dagnino-Subiabre, A., Silva, J., & Lopez, V. (2023). Impact of acute psychosocial stress on attentional control in humans. A study of evoked potentials and pupillary response. Neurobiology of Stress, 25, 100551. 10.1016/j.ynstr.2023.100551

Rudebeck, P. H., & Rich, E. L. (2018). Primer: The Orbitofrontal Cortex. Current Biology : CB, 28(18), R1083–R1088. 10.1016/j.cub.2018.07.018

Salsman, J. M., Butt, Z., Pilkonis, P. A., Cyranowski, J. M., Zill, N., Hendrie, H. C., Kupst, M. J., Kelly, M. A. R., Bode, R. K., Choi, S. W., Lai, J.-S., Griffith, J. W., Stoney, C. M., Brouwers, P., Knox, S. S., & Cella, D. (2013). Emotion assessment using the NIH Toolbox. Neurology, 80(11_supplement_3), S76–S86. 10.1212/WNL.0b013e3182872e11

Sandi, C. (2013). Stress and cognition. Wiley Interdisciplinary Reviews. Cognitive Science, 4(3), 245–261. 10.1002/wcs.1222

Shackman, A. J., Maxwell, J. S., McMenamin, B. W., Greischar, L. L., & Davidson, R. J. (2011). Stress Potentiates Early and Attenuates Late Stages of Visual Processing. The Journal of Neuroscience, 31(3), 1156–1161. 10.1523/JNEUROSCI.3384-10.2011

Sonkusare, S., Breakspear, M., & Guo, C. (2019). Naturalistic Stimuli in Neuroscience: Critically Acclaimed. Trends in Cognitive Sciences, 23(8), 699–714. 10.1016/j.tics.2019.05.004

Tiferet-Dweck, C., Hensel, M., Kirschbaum, C., Tzelgov, J., Friedman, A., & Salti, M. (2016). Acute stress and perceptual load consume the same attentional resources: A behavioral-ERP study. PLoS ONE, 11(5). 10.1371/journal.pone.0154622

Viviani, R. (2014). Neural Correlates of Emotion Regulation in the Ventral Prefrontal Cortex and the Encoding of Subjective Value and Economic Utility. Frontiers in Psychiatry, 5. https://www.frontiersin.org/articles/10.3389/fpsyt.2014.00123

Wang, J., Yu, L., Ding, F., & Qi, C. (2023). Effect of Acute Psychological Stress on Speed Perception: An Event-Related Potential Study. Brain Sciences, 13(3), Article 3. 10.3390/brainsci13030423

Wearne, T. A., Lucien, A., Trimmer, E. M., Logan, J. A., Rushby, JacquelineA., Wilson, E., Filipčíková, M., & McDonald, S. (2019). Anxiety sensitivity moderates the subjective experience but not the physiological response to psychosocial stress. International Journal of Psychophysiology, 141, 76–83. 10.1016/j.ijpsycho.2019.04.012

Welborn, B. L., Papademetris, X., Reis, D. L., Rajeevan, N., Bloise, S. M., & Gray, J. R. (2009). Variation in orbitofrontal cortex volume: Relation to sex, emotion regulation and affect. Social Cognitive and Affective Neuroscience, 4(4), 328–339. 10.1093/scan/nsp028

Wu, Y., Wang, J., Zhang, Y., Zheng, D., Zhang, J., Rong, M., Wu, H., Wang, Y., Zhou, K., & Jiang, T. (2016). The Neuroanatomical Basis for Posterior Superior Parietal Lobule Control Lateralization of Visuospatial Attention. Frontiers in Neuroanatomy, 10. 10.3389/fnana.2016.00032

Xu, C., Xu, Y., Xu, S., Zhang, Q., Liu, X., Shao, Y., Xu, X., Peng, L., & Li, M. (2020). Cognitive Reappraisal and the Association Between Perceived Stress and Anxiety Symptoms in COVID-19 Isolated People. Frontiers in Psychiatry, 11. https://www.frontiersin.org/articles/10.3389/fpsyt.2020.00858

Yang, M., Tsai, S.-J., & Li, C.-S. R. (2020). Concurrent amygdalar and ventromedial prefrontal cortical responses during emotion processing: A meta-analysis of the effects of valence of emotion and passive exposure versus active regulation. Brain Structure and Function, 225(1), 345–363. 10.1007/s00429-019-02007-3

Yaribeygi, H., Panahi, Y., Sahraei, H., Johnston, T. P., & Sahebkar, A. (2017). The impact of stress on body function: A review. EXCLI Journal, 16, 1057. 10.17179/excli2017-480

Zhu, X., Yan, W., Lin, X., Que, J., Huang, Y., Zheng, H., Liu, L., Deng, J., Lu, L., & Chang, S. (2022). The effect of perceived stress on cognition is mediated by personality and the underlying neural mechanism. Translational Psychiatry, 12, 199. 10.1038/s41398-022-01929-7

