## Supplementary Materials for "Lower perceived stress enhances neural synchrony in perceptual and attentional cortices during naturalistic processing"

**Supplementary Materials A**

**Table S1**

*Pearson Correlations and P-values Among Model Variables*

| Measures | Perceived Stress (Pearson Correlation) | Sig. (2-tailed) |
| --- | --- | --- |
| Social Satisfaction | -.372* | .0012 |
| Sorting Working Memory Test | -0.161 | 0.141 |
| Pattern Comparison Processing Speed Test | -.321 | .781 |
| Picture Sequence Memory Test | .061 | .624 |
| Psychological Well Being | -.479* | .001 |
| Flanker Inhibitory Control and Attention Test | -1.11 | .312 |
| Dimensional Change Card Sort Test | .111 | .382 |
| Oral Symbol Digit Test | -.121 | .342 |
| Auditory Verbal Learning Test | .062 | .552 |
| Positive Affect | -.532* | .0014 |
| General Life Satisfaction | -.541* | .0013 |
| Meaning and Purpose | -.422* | .0012 |
| Emotional Support | -.142 | .212 |
| Instrumental Support | -.221* | .041 |
| Friendship | -.401* | .0016 |
| Loneliness | .321* | .023 |
| Perceived Rejection | .121 | .291 |
| Perceived Hostility | .161 | .154 |
| Self-Efficacy | -.491* | .0015 |
| Fear-Affect | .063 | .584 |
| Fear-Somatic Arousal | .062 | .563 |
| Sadness | .011 | .994 |
| Anger-Affect | .342* | .031 |
| Anger-Hostility | .642* | .001 |
| Anger-Physical Aggression | .031 | .811 |
| Age | .623 | .611 |

***Note.*** ** Denotes significance at p < 0.01, ** denotes significance at the Bonferroni corrected value of p < 0.002.  No correlations were significant at this threshold.*

**Low-Stress Average Cluster Table**

**Table S2**

*Coordinates and cluster sizes for ISC found within the low-low PSS group average contrast. Clusters with five or more voxels shown.*

| **Anatomical Location** | **Hemisphere** | **# of Voxels** | **MAX *r*** | **MAX x** | **MAX y** | **MAX z** |
| --- | --- | --- | --- | --- | --- | --- |
| Intracalcarine Cortex | Left | 325 | 0.242 | -10.5 | -82.5 | 7.5 |
| Superior Temporal Gyrus, anterior division | Left | 300 | 0.182 | -61.5 | -4.5 | -4.5 |
| Lateral Occipital Cortex, inferior division | Right | 298 | 0.199 | 52.5 | -64.5 | 7.5 |
| Superior Temporal Gyrus, anterior division | Right | 290 | 0.187 | 60.5 | 3.5 | -5.5 |
| Lateral Occipital Cortex, inferior division | Left | 142 | 0.241 | -49.5 | -70.5 | 10.5 |
| Occipital Fusiform Gyrus | Right | 135 | 0.147 | 22.5 | -70.5 | -7.5 |
| Intracalcarine Cortex | Right | 129 | 0.321 | 10.5 | 80.5 | -7.5 |
| Temporal Occipital Fusiform Cortex | Left | 100 | 0.126 | -28.5 | -55.5 | -7.5 |
| Planum Temporale | Left | 99 | 0.327 | -61.5 | -16.5 | 7.5 |
| Central Opercular Cortex | Right | 52 | 0.247 | 58.5 | -10.5 | 7.5 |
| Cingulate Gyrus, posterior division | Right | 23 | 0.0572 | 13.5 | -22.5 | 40.5 |
| Precentral Gyrus | Right | 22 | 0.0275 | 37.5 | -22.5 | 67.5 |
| Lateral Occipital Cortex, inferior division | Left | 20 | 0.0907 | -49.5 | -67.5 | -10.5 |
| Precentral Gyrus | Right | 19 | 0.128 | 58.5 | -1.5 | 46.5 |
| Temporal Parietal Junction | Right | 19 | 6.32 | 46.5 | 43.5 | -25.5 |
| Juxtapositional Lobule Cortex | Left | 19 | 0.0532 | -1.5 | 1.5 | 67.5 |
| Frontal Pole | Right | 18 | 0.047 | 49.5 | 40.5 | 4.5 |
| Precentral Gyrus | Left | 16 | 0.123 | -52.5 | -4.5 | 52.5 |
| Orbital Frontal Cortex | Right | 16 | 0.0265 | 31.5 | 25.5 | -4.5 |
| Precuneus Cortex | Right | 15 | 0.0738 | 9.5 | 45.5 | -54.5 |
| Precentral Gyrus | Left | 15 | 0.0267 | -43.5 | -13.5 | 58.5 |
| Superior Frontal Gyrus | Right | 14 | 0.0332 | 28.5 | 22.5 | 58.5 |
| Superior Frontal Gyrus | Left | 13 | 0.0296 | -4.5 | 13.5 | 55.5 |
| Lateral Occipital Cortex, superior division | Right | 11 | 0.0779 | 13.5 | -58.5 | 64.5 |
| Occipital Pole | Left | 9 | 0.17 | -13.5 | -91.5 | 31.5 |
| Frontal Pole | Left | 9 | 0.039 | -1.5 | 61.5 | 25.5 |
| Precuneus Cortex | Left | 8 | 0.0708 | -10.5 | -46.5 | 55.5 |
| Occipital Pole | Right | 8 | 0.131 | 10.5 | -94.5 | 19.5 |
| Precentral Gyrus | Left | 7 | 0.0238 | -7.5 | -28.5 | 79.5 |
| Lateral Occipital Cortex, superior division | Right | 6 | 0.0569 | 34.5 | -58.5 | 61.5 |
| Superior Parietal Lobule | Right | 6 | 0.0735 | 30.5 | -43.5 | 58.5 |
| Lateral Occipital Cortex, superior division | Left | 6 | 0.0631 | -55.5 | -61.5 | 25.5 |
| Middle Frontal Gyrus | Right | 5 | 0.0554 | 52.5 | 25.5 | 25.5 |
| Lateral Occipital Cortex, superior division | Left | 5 | 0.0794 | -31.5 | -61.5 | 61.5 |
| Superior Parietal Lobule | Left | 5 | 0.0716 | -32.5 | -42.5 | 57.5 |
| Lateral Occipital Cortex, superior division | Left | 5 | 0.0611 | -22.5 | -64.5 | 64.5 |

**Table S3**

*Coordinates and cluster sizes for ISC found within the high-high PSS group average contrast. Clusters with five or more voxels shown.*

| **Anatomical Location** | **Hemisphere** | **# of Voxels** | **MAX *r*** | **MAX x** | **MAX y** | **MAX z** |
| --- | --- | --- | --- | --- | --- | --- |
| Lateral Occipital Cortex, superior division | Left | 78 | 0.00835 | -31.5 | -85.5 | 7.5 |
| Intracalcarine Cortex | Right | 75 | 0.00377 | 16.5 | -73.5 | 13.5 |
| Lateral Occipital Cortex, superior division | Right | 74 | 0.00347 | 46.5 | -79.5 | 16.5 |
| Superior Temporal Gyrus, posterior division | Right | 71 | 0.00599 | 67.5 | -19.5 | -1.5 |
| Intracalcarine Cortex | Left | 70 | 0.00836 | -10.5 | -79.5 | 7.5 |
| Planum Temporale | Right | 68 | 0.355 | 61.5 | -7.5 | 4.5 |
| Planum Temporale | Left | 63 | 0.361 | -61.5 | -13.5 | 7.5 |
| Superior Temporal Gyrus, posterior division | Left | 52 | 0.004 | -52.5 | -37.5 | 1.5 |
| Parahippocampal Gyrus | Right | 48 | 0.048 | 1.8 | -25.5 | 41.4 |
| Parahippocampal Gyrus | Left | 48 | 0.0582 | -1.8 | 25.5 | 43.1 |
| Precentral Gyrus | Right | 33 | 0.062 | 28.5 | -7.5 | -61.5 |
| Supramarginal Gyrus, posterior division | Right | 32 | 0.0438 | 61.5 | -43.5 | 34.5 |
| Juxtapositional Lobule Cortex | Left | 19 | 0.0532 | -1.5 | 1.5 | 67.5 |
| Frontal Pole | Right | 18 | 0.047 | 49.5 | 40.5 | 4.5 |
| Inferior Frontal Gyrus, pars opercularis | Left | 13 | 0.00279 | -52.5 | 19.5 | 25.5 |
| Inferior Frontal Gyrus, temporooccipital part | Right | 11 | 0.00153 | 58.5 | -55.5 | -10.5 |
| Superior Parietal Lobule | Left | 9 | 0.0736 | 31.5 | -43.5 | 58.5 |
| Occipital Pole | Left | 8 | 0.131 | -10.5 | 94.5 | -19.5 |
| Precentral Gyrus | Left | 7 | 0.0238 | -7.5 | -28.5 | 79.5 |
| Lateral Occipital Cortex, superior division | Right | 6 | 0.0569 | 34.5 | -58.5 | 61.5 |
| Lateral Occipital Cortex, superior division | Left | 6 | 0.0631 | -55.5 | -61.5 | 25.5 |
| Lateral Occipital Cortex, superior division | Left | 6 | 0.0793 | -30.5 | -61.5 | 61.5 |
| Superior Parietal Lobule | Left | 5 | 0.0716 | -31.5 | -43.5 | 56.5 |
| Lateral Occipital Cortex, superior division | Left | 5 | 0.0611 | -22.5 | -64.5 | 64.5 |

**Tables S4**

*Coordinates and cluster sizes for ISC found within the low-low PSS group compared to the high-high PSS group. Clusters with five or more voxels shown.*

| **Anatomical Location** | **Hemisphere** | **# of Voxels** | **MAX *r*** | **MAX x** | **MAX y** | **MAX z** |
| --- | --- | --- | --- | --- | --- | --- |
| Lateral Occipital Cortex | Right | 18 | 0.41 | 19.9 | 61.5 | -9.5 |
| Lateral Occipital Cortex | Left | 13 | 0.41 | -19.7 | -60.1 | 9.5 |
| Superior Parietal Lobule | Right | 12 | 0.0736 | 29.5 | 41.5 | -56.5 |
| Superior Temporal Gyrus | Right | 11 | 0.176 | 62.5 | 18.5 | -7.5 |
| Orbital Frontal Cortex | Right | 7 | 0.0265 | 31.5 | 25.5 | -4.5 |
| Occipital Pole | Right | 6 | 0.121 | 9.5 | -94.5 | 18.5 |
| Superior Temporal Gyrus | Left | 5 | 0.176 | -64.5 | -19.5 | 7.5 |
| Intracalcarine Cortex | Right | 5 | 0.120 | 15.5 | 103.8 | -60.5 |
| Intracalcarine Cortex | Left | 5 | 0.0745 | -13.5 | -94.5 | 34.5 |
| Occipital Pole | Left | 5 | 0.131 | -11.5 | 93.5 | -18.5 |
| Superior Parietal Lobule | Left | 5 | 0.0736 | -31.5 | -43.5 | 58.5 |
